# Insights from the first genome assembly of Onion (*Allium cepa*)

**DOI:** 10.1101/2021.03.05.434149

**Authors:** Richard Finkers, Martijn van Kaauwen, Kai Ament, Karin Burger-Meijer, Raymond Egging, Henk Huits, Linda Kodde, Laurens Kroon, Masayoshi Shigyo, Shusei Sato, Ben Vosman, Wilbert van Workum, Olga Scholten

## Abstract

Onion is an important vegetable crop with an estimated genome size of 16Gb. We describe the *de novo* assembly and *ab initio* annotation of the genome of a doubled haploid onion line DHCU066619, which resulted in a final assembly of 14.9 Gb with a N50 of 461 Kb. Of this, 2.2 Gb was ordered into 8 pseudomolecules using five genetic linkage maps. The remainder of the genome is available in 89.8 K scaffolds. Only 72.4% of the genome could be identified as repetitive sequences and consist, to a large extent, of (retro) transposons. In addition, an estimated 20% of the putative (retro) transposons had accumulated a large number of mutations, hampering their identification, but facilitating their assembly. These elements are probably already quite old. The *ab initio* gene prediction indicated 540,925 putative gene models, which is far more than expected, possibly due to the presence of pseudogenes. Of these models, 86,073 showed similarity to published proteins (UNIPROT). No gene rich regions were found, genes are uniformly distributed over the genome. Analysis of synteny with *A. sativum* (garlic) showed collinearity but also major rearrangements between both species. This assembly is the first high-quality genome sequence available for the study of onion and will be a valuable resource for further research.

## Introduction

More than just a tasty culinary sensation, onion (*Allium cepa* L.) is one of the most important vegetable crops worldwide. In terms of global production value, onion ranks second after tomato (http://www.fao.org/faostat/en/#home). Onion is a diploid (2*n* = 2*x* = 16) species with a genome size of approx. 16,400 Mb/1C (Van’T Hof 1965; Arumuganathan and Earle 1991; Ricroch *et al.* 2005), the largest of all cultivated diploid crops and of a size comparable to the allo-hexaploid bread wheat (Brenchley *et al.* 2012; Marcussen *et al.* 2014). A large genome size is often associated with repeat accumulation (Kelly and Leitch 2011). The C_o_T reannealing kinetics indicate that about 40% of the onion genome is highly repetitive (>1000x copies) and 40% has 100-1000 copies and is thus middle to low repetitive (Stack and Comings 1979). Overall, at least 95% of the *A. cepa* genome consists of repetitive sequences (Flavell *et al.* 1974), most of which are dispersed repeats (Shibata and Hizume 2002) and LTR retrotransposons of the Ty1/copia and Ty3/gypsy type (Pearce *et al.* 1996; Kumar *et al.* 1997; Pich and Schubert 1998; Vitte *et al.* 2013). Due to the size of the genome and the repetitive nature, developing an onion reference genome assembly is challenging (Havey and McCallum 2012).

For onion, molecular breeding strategies are currently limited to the use of molecular markers and genetic linkage maps (Martin *et al.* 2005; McCallum *et al.* 2012; Baldwin *et al.* 2012; Scholten *et al.* 2016). Knowledge of the genome of onion and related species is scarce compared to other crop plants, with only transcriptome sequences available (Kim *et al.* 2014; Kamenetsky *et al.* 2015; Sohn *et al.* 2016; Abdelrahman *et al.* 2017). While the availability of a reference genome has greatly stimulated research and led to accelerated breeding in many other crops (Kuhl *et al.* 2005; Vitte *et al.* 2013; Sun *et al.* 2020), onion has not yet had this benefit. Garlic (*Allium sativum*; 16.2 Gb) and asparagus (*Asparagus officinalis* L.; 1.1 Gb) are the most closely related species with reference genomes available (Harkess *et al.* 2017; Sun *et al.* 2020). Though, useful for gene discovery, the lack of insight into detailed syntenic relationship between these crops and onion is still limiting the utilization in onion (breeding) research.

In this paper, we describe the first *de novo* assembly of the genome of a doubled haploid *A. cepa* accession through a combination of strategies and the development of an initial set of pseudomolecules. Synteny between onion and garlic was studied and provides a first insight into the similarities and differences in genome organization. This onion assembly will be an important tool, facilitating onion breeding and research.

## Methods & Materials

### Plant material

Seeds of the doubled haploid (DH) *Allium cepa* line DHCU066619 (Hyde *et al.* 2012) were kindly provided by Dr. M. Mutschler (Cornell University). This DH accession was selected for whole-genome shotgun sequencing (WGS), as it is a homozygous, vigorously growing genotype.

### DNA and RNA isolation

Two grams of leaf tips from young and rapid-growing onion plants were harvested and pooled for nuclei DNA isolation according to the protocol described by Bernatzky and Tanksley (1986).

For RNA extraction, tissues from bulb and basal plate, as well as leaf and flowers from mature plants were harvested, frozen in liquid nitrogen and grinded. Approximately 100 mg of each tissue was transferred to a 2ml screwcap tube, followed by adding 800 μl of Trizol and vortexing for 1 minute. Subsequently 160 μl of Chloroform was added, and gently mixed for 15 seconds. After incubating the samples for 3 minutes at RT, they were centrifuged at 10,000 rpm in an Eppendorf centrifuge for 3 minutes, followed by transfer of the water-phase to a clean tube. 350 μl of RLT buffer (Qiagen) containing 10 μg/μl β-mercaptoethanol, was added to each 100 μl of water-phase, followed by a RNeasy (Qiagen) column extraction.

### Sequencing methods, preparation details and data processing

Three TruSeq sequencing libraries (with median insert sizes of 230, 350, and 500) were made according to the manufacturer’s recommendations and sequenced during a single Illumina^®^ HiSeq 2500 run (across 16 lanes) by GenomeScan (Leiden, The Netherlands). Quality of the data was assessed using fastQC (https://www.bioinformatics.babraham.ac.uk/projects/fastqc/) and by evaluating the k-mer profile using JELLYFISH (Marçais and Kingsford 2011).

PacBio Sample preparation was performed according to the PacBio protocol “*20 kb Template Preparation Using BluePippin™ Size-Selection System”*. In short, 8 μg of sample was fragmented using a Covaris g-Tube at 4800 rpm for 1 minute. The PacBio SMRTbell™ Template Prep Kit 1.0 was used for the DNA library preparation of the primer annealed SMRTbells. The SMRTbells were size selected using the BluePippin set for 10kb-50kb long reads. The PacBio DNA/Polymerase Binding Kit P6 was used to bind prepared SMRTbell libraries to the DNA polymerase in preparation for sequencing on the PacBio *RS* II. The complex of polymerase bound SMRTbell was mixed with long-term storage buffer. Two batches of stored complex were made. Prior to sequencing the SMRTbell complex was incubated with PacBio MagBeads. Sequencing was performed with the PacBio DNA Sequencing Kit 4.0 chemistry. For the sequencing run MagBead loading and Stage Start were enabled. Twenty sequencing runs, totaling 138 SMRTcells, were performed with a 360-minute movie time per SMRTcell. The sequence runs were performed on the PacBio RS II sequencer and primary analysis was performed with the SMRT Analysis server version 2.3.0 (GenomeScan, The Netherlands).

### Genome Assembly

Illumina reads were assembled using the MaSuRCA 2.3.1 assembly pipeline according to the author’s recommendations (Zimin *et al.* 2013). The PacBio reads were assembled with the Illumina assembly as a backbone using DBG2OLC (Ye *et al.* 2016a) and varying settings for kmerSize, KmerCov, MinOverlap and AdaptTH (File S2; File S3). Final DBG2OLC assembly was performed with KmerSize=21, KmerCov =2, MinOverlap=20 and AdaptThr=0.05.

### Genome Scaffolding

#### Dovetail scaffolding

Five grams of young leaf tissue (lyophilized) was sent to Dovetail Genomics (USA) for High Molecular Weight (HMW) DNA isolation, together with the PacBio/Illumina hybrid assembly, the unplaced Illumina scaffolds > 3Kb and a published onion chloroplast sequence (von Kohn *et al.* 2013) to produce a scaffolded assembly according to the Chicago protocol (Putnam *et al.* 2016). For this, the generated DOVETAIL sequencing libraries were analyzed with a modified version of the HiRise algorithm (Dovetail Genomics, USA) to accommodate the large genome size.

#### Genetic marker-based scaffolding

The dovetail scaffolded genome assembly was anchored into pseudomolecules with Allmaps (Tang *et al.* 2015) using five previously published genetic linkage maps (Shigyo; Duangjit *et al.* 2013; Scholten *et al.* 2016; Choi *et al.* 2020).

### Genome Annotation

#### Completeness

Completeness of the onion genome assembly was assessed by BUSCO v4.1.1; database version embryophyte_obd10 2019-11-27 (Simao *et al.* 2015).

#### Repeat annotation

To structurally annotate repeat sequences in the *A. cepa* genome *de novo*, RepeatMasker was applied using the REPBASE v20.5 library. *Ab initio* prediction of repeats was performed using the TEdenovo pipeline of REPET v2.5 (Flutre *et al.* 2011), with default parameters, utilizing NCBI-Blast+. To reduce computational time needed to execute the TEdenovo pipeline, 0.52 Gb of the 14.4 Gb (3.6%) of the onion assembly was selected at random. Grouper, Recon and Piler steps were invoked both with and without structural detection. Repbase v20.05 and Pfam27.0 HMM profiles were used to annotate repeats identified in onion by the pipeline. The output of the TEdenovo pipeline was subsequently used as the reference library to run the TEannot pipeline on randomized chucks of the whole assembly using default parameters and exported to GFF3 format.

#### *Ab initio* gene prediction

RNA from four tissues (bulb, basal plate, leaf, and flower) was isolated and sequenced by GenomeScan (Leiden, The Netherlands) using an Illumina HiSeq 2500 sequencer according to the manufacture’s recommendations. BRAKER1 (Hoff *et al.* 2015) was used for unsupervised training of Augustus 3.2.2 (Stanke *et al.* 2008), using splice junctions identified from mapping RNAseq reads with STAR mapper (Dobin *et al.* 2013) of all four tissues combined. Subsequently, the training parameters from BRAKER1 were used to annotate the masked genome sequence.

#### Functional annotation / Blast2go interpro

Blast2Go (Gotz *et al.* 2008) was used for functional annotation of the predicted protein models with the default settings for the mapping and annotation step. The initial blastp 2.6.0+ step was performed against the Swissport database (version 4 Oct. 2017) with an e-value cut-off of 1,0E−3, word size of 6, Low Complexity filter on true, and a maximum of 20 blast hits. InterProScan v5.26 (Jones *et al.* 2014) including panther 12.0 libraries was used to identify protein domains within the predicted protein sets.

### Synteny analysis

EST based markers from Shigyo et al. (2020) were blasted to the onion genome and to the garlic genome. 3904 markers had a hit to both genome sequences, of which 781 positioned at unanchored onion scaffolds, leaving 3123 markers positioned at the onion pseudomolecules. Based on the top hits, the physical positions of markers with a match to both genomes were plotted against each other in a XY plot using the python matplotlib and seaborn libraries.

### Data availability

The onion DH is deposited to the U.S. National Plant Germplasm System and is available under the number: G 32985. The final assembly and annotation files are available on www.oniongenome.wur.nl for download and in a Genome browser (Skinner *et al.* 2009). All data, including the genome sequence and raw sequencing reads have been deposited to EBI under the BioProject ID PRJEB29505.

## Results and Discussion

### Genome assembly

The doubled haploid *Allium cepa* accession DHCU066619 (Hyde *et al.* 2012) was selected for whole-genome shotgun sequencing (WGS), to facilitate genome assembly especially of a large genome like onion. Three small insert libraries were used for Illumina HiSeq 2500 sequencing (Table S1) and yielded 769 Gb sequence data. Analysis of the sequence libraries resulted in ~450 G *k*-mers (k=31), and an estimated genome size of approx. 13.6 Gb. Of this, approx. 7.4 Gb (53.8%) is single copy based on *k*-mer statistics, indicating that an initial assembly from Illumina reads is feasible. The MaSurCa based assembly resulted in 10.8 Gb in 6.2M contigs with a contig N50 of 2.7Kb (Table 1). This assembly was further scaffolded using 18.1M PacBio RS II reads >1KB with DBG2OLC. This resulted in an assembly of 14.6 Gb in 316K contigs and a contig N50 size of 59Kb (Table 1). In the last step, the hybrid Illumina/PacBio assembly was further improved using Dovetail Chicago and subsequent HiRise scaffolding which indicated that there were no mis-assemblies in the Illumina/PacBio hybrid contigs, showing the high quality of this initial assembly. The combination of these three technologies resulted in an assembly of 14.9 Gb in 92.9K scaffolds with a scaffold N50 size of 436Kb. With an estimated genome size of 16,400 Gb/1C, we managed to assemble ~91% of the onion genome. As our assembly is mostly based on short read Illumina sequencing, with limited data from third generation PacBio long reads, we hypothesize that the most complex highly repetitive regions are missing from our assembly.

**Table 1.** Statistics of the different steps in the onion genome assembly process. ADD ABBREVIATIONS IN DESCRIPTION OF THE TABLE. Add comments about Not Determined fields. * Gene statistics based on 86k models with blast support

|  | Illumina | Illumina & PacBio hybrid | Illumina, PacBio & Dovetail | Illumina, PacBio, Dovetail & Allmaps – A. cepa v1.0 |
| --- | --- | --- | --- | --- |
| Total assembly size (GB) | 10.80 | 14.59 | 14.94 | 14.94 |
| N50 (bp) | 2,776 | 59,009 | 436,203 | 460,718 |
| L50 | 91,028,083 | 76,344 | 9,266 | 6,712 |
| NR of Contigs (K) | 6,200 | 316 | - | - |
| NR of Scaffolds (K) | - | - | 92.9 | 89.8 |
| NR of Scaffolds incorporated in Pseudomolecules (K) | - | - | - | 3.0 |
| Assembly size in Pseudomolecules (Gb) | - | - | - | 2.2 |
| G + C content (%) | 33.2 | 33.5 | 33.5 | 33.5 |
| Estimated TE content (%) | - | - | - | 72.4 |
| Number of gene models | - | - | - | 540,925 |
| Number of gene models with >90% RNAseq support | - | - | - | 47,066 |
| Median / Average /max Exon length (bp)* | - | - | - | 137 / 202/ 14,836 |
| Median / Average / Max intron length (bp)* | - | - | - | 178 / 1,035 / 213,043 |
| N’S per 100 Kbp | 0 | 0 | 2,344 | 2,346 |

### Anchoring scaffolds into pseudomolecules using multiple EST based genetic maps

To further organize our genome assembly, we used three intraspecific genetic linkage maps (Shigyo; Duangjit *et al.* 2013; Choi *et al.* 2020) and two interspecific genetic linkage maps (Scholten *et al.* 2016) to anchor scaffolds into pseudomolecules using AllMaps (Tang *et al.* 2015). Except for the GBS based markers from the study of Choi *et al*. (2020), all markers were developed from transcriptome sequencing. With this approach, we were able to anchor 3,028 scaffolds (3.2% of the total number of scaffolds) into eight pseudomolecules, with an overall length of 2.2Gb (14.9% of the assembly size; File S6). The pseudomolecules were named according to the Linkage group (LG) assignments published by van Heusden et al. (2000); which follows the system of chromosome nomenclature for *A. cepa* according to de Vries (1990). A subset of 635 scaffolds (0.6Gb) could be oriented according to the genetic map order. The other 1,910 scaffolds could not as they were anchored based on one marker only. Overall agreement between the scaffolds and genetic positions was good (File S6) with absolute Spearman correlation coefficients over the five maps ranging between 0.69 (CCxRR) and 0.95 (GBS). Notably, the two interspecific maps (CCxRR, which is an F2 of onion x *A. roylei*; avg ρ=0.69 and CCxRF, which is a cross between onion and the interspecific F1 hybrid between *A. roylei* and *A. fistulosum*; avg ρ=0.74; (Scholten *et al.* 2016)) show a lower overall spearman correlation coefficient than the intraspecific maps, which is to be expected as, for example, structural variation such as rearrangements and inversions will break collinearity. Disruption of collinearity in homeologous chromosomes in Allium species has been reported, for example in chromosome 4 (Khrustaleva *et al.* 2019a). Omitting the two interspecific maps from the Allmaps procedure did not significantly impact the Allmaps scaffolding (data not shown) and therefore we included these maps in the result. While the overall order of chrLG1 between the five maps was very consistent (spearman correlation coefficient of 0.95), chrLG7 showed the lowest correlation (spearman rho coefficient of 0.69), because of a low correlation coefficient of the two interspecific maps (CCxRR; ρ=0.32 and CCxRF; ρ=0.54), indicating that there are different levels in overall synteny per chromosome between the different species. Not all maps were informative in the Allmaps scaffolding. Markers of the CCxRF map did not contribute to the scaffolding of chrLG3 while markers of the GBS map did not contribute to the scaffolding of chrLG3, chrLG5 and chrLG8. ALLMAPS has implemented an algorithm to minimize ambiguity (Tang *et al.* 2015), and the GBS marker based sequences showed similarity to more than one scaffold, which were assigned to different linkage group, and were therefore skipped for the final scaffolding. Overall, the developed pseudomolecules will be very useful in developing additional genetic markers, fine mapping QTL regions and/or candidate gene mining.

### Genome annotation

The completeness of the Dovetail scaffolded *de novo* genome assembly was evaluated using BUSCO genes (embryophyta_odb10; 2020-09-10; File S4) and resulted in a completeness score of 87.7%. Other large genome assemblies, such as the *loblolly pine* v1.01 genome has an CEGMA completeness of 91% (Neale *et al.* 2014) and the *Allium sativum* v1.0 genome has an CEGMA completeness score of 92.7% and a BUSCO completeness score of 88.7% (Sun *et al.* 2020). Of the BUSCO genes, 3.3% were labeled as fragmented, which could be because the length of the gene model did not fall within the expected length distribution of the chosen BUSCO profile. Also technical limitation of the algorithm might increase proportions of fragmented and missing BUSCOs, especially in large genomes (Simao *et al.* 2015). Still, the slightly lower completeness scores in onion suggests that we miss genes in our assembly.

Initial repeat masking of the genome with repeatmasker, using the REPBASE v20.5 database, resulted in 15.1 % of the genome to be annotated as repetitive (Table S5), which is far below the expected 95% moderate to high repetitive regions (Flavell *et al.* 1974) though comparable to homology based repeat annotation in other large genomes, such as *loblolly pine* (Wegrzyn *et al.* 2013). This indicates that most repeats in the genome are not recognizable anymore as repeats, have accumulated mutations, and are probably (very) old. This observation agrees with the observed Kmer statistics, which suggests that 53,8% of the genome is single copy and the observations by Jakše *et al.* (2008) that onion BAC sequences contain > 50 % sequences that are like transposons, many of which are degraded. Using *de novo* repeat annotation strategies, genomes comparable in size, such as garlic (Sun *et al.* 2020), bread wheat (Mayer *et al.* 2014) and loblolly pine (Wegrzyn *et al.* 2014) were annotated as containing approx. 91%, 80%, and 82% repetitive DNA sequences, respectively. Therefore, we used the ‘all vs all comparison methods’, as implemented in the REPET pipeline (Flutre 2009) to develop an onion specific repeat database. Using this *de novo* developed repeat database, 72.4% of the genome sequence could be classified as repetitive and was subsequently masked for downstream analysis. Still, this percentage is lower than the expected 95%, suggesting that the remaining ~20% of the “repetitive” sequences were too diverged to be recognized.

Long terminal repeat (LTR) retrotransposons are the major contributors to the size of the onion genome (table S5), which is in line with previous studies (Vitte *et al.* 2013). The REPBASE annotation indicates that the majority of the young LTRs are of the Gypsy type; followed by Copia. This has also been observed in other large plant genomes such as spruce (Nystedt *et al.* 2013), garlic (Sun *et al.* 2020) and wheat (Appels *et al.* 2018). LTR retrotransposons have been described in relation to genome size increase (Kumar and Bennetzen 1999) and have been suggested to play a significant role in adaptive response of the genome to environmental challenges (McClintock 1984).

*Ab initio* prediction of gene models on the repeat masked genome sequence with Augustus resulted in the identification of 540,925 gene models, from which 47,066 showed > 90% coverage with reads from the RNAseq dataset (Table 1). The number of gene models is way beyond the average number of genes of 36,795 reported for plant genomes (Ramírez-Sánchez *et al.* 2016). A proper annotation to identify only functional genes would require extensive manual curation (Hosmani *et al.* 2019b; Tello-Ruiz *et al.* 2019; Athanasouli *et al.* 2020). We decided to make the extended set available for the community, rather than restricting ourselves to making only models available with additional RNAseq support. The abundance of gene models may most likely be explained by the presence of pseudogenes (Zhang and Gerstein 2004; Xiao *et al.* 2016). Pseudogenes are non-functional copies of genes that were once active in the ancestral genome. For example, in Arabidopsis 924 pseudogenes are known (Xiao *et al.* 2016) while in wheat 288,839 pseudogenes were identified (Appels *et al.* 2018). A pseudogene still has characteristics of a gene and will be detected using an *ab initio* gene model prediction algorithm, but not be annotated by blast against curated protein databases, such as TrEMBL, due to partial matches. Blast analysis against TrEMBL resulted in hits for 86,073 gene models (15.9% of the *ab initio* predicted models). Using Blast2Go (Conesa *et al.* 2005), 88,259 gene models were functionally annotated, of which 49,918 models were annotated using data from both Blast and InterPro. Of these, 17,457 models were annotated exclusively by InterPro, while 20,884 models had a blast hit only. For subsequent analysis, we focus on the subset of 86,073 models with similarity to genes from TrEMBL, of which 25,344 showed > 90% coverage with reads from the RNAseq dataset. The average coding sequence length of onion genes is 879 bp and is spread over 4.4 exons. This is shorter than the average plant gene length of 1,308 bp (Ramírez-Sánchez *et al.* 2016), though still within the observed variation and larger than the average gene length of 797 bp reported for garlic (Sun *et al.* 2020). Average intron length is 1,035 bp (Table 1), though the largest predicted intron is 213 Kb. Although a positive relationship between intron size and genome size has been observed (Stival Sena *et al.* 2014), due to large variations that occur in intron sizes, it is not a good predictor for genome size (Wendel *et al.* 2002). With a median and average intron size of 178 and 1,035 bp respectively, the majority of introns in onion is short, while a limited number of introns is (very) long. This is probably because of the energy required for transcribing the long genes.

### Organization of the onion gene space

In plants, we have seen two scenario’s for the distribution of genes: genes mainly located in actively recombining euchromatin regions, while large non-recombining regions (centromeres) have a low gene content, such as in tomato (Hosmani *et al.* 2019a), or genes equally distributed over the genome, such as in garlic (Sun *et al.* 2020). If onion would show a similar pattern as tomato, then most gene models will be included in the current set of pseudomolecules. Based on the set of 86K models with a match to the TrEMBL database, we calculated a rate of 8,108 and 5,352 gene models/Gb for anchored and unanchored scaffolds, respectively. This shows that gene density in the anchored scaffolds is approximately 1.5x higher than in the unanchored scaffolds. In tomato, we calculated gene density in euchromatin and heterochromatin (Sim *et al.* 2012; Víquez-Zamora *et al.* 2014) to be 97,715 and 20,010 gene models/Gb respectively, a difference of approximately 4.9x. Interpretation of this data and the meaning for onion must be treated with caution as it is influenced by two factors; I) the difference in genome size between onion (16Gb) and tomato (850Mb) and II) the fact that only 2.2Gb (out of 14.9Gb) of the onion scaffolds are yet organized into pseudomolecules. Having said that, like Jakse et al. (2008) who previously sequenced two onion BACs, we hypothesize that genes in onion are more equally distributed over the genome, and reside in an ocean of repetitive elements, e as the ratio between scaffolds incorporated in pseudomolecules and the scaffolds not incorporated in pseudomolecules is much closer to 1, than the ratio observed in tomato. This hypothesis is further supported by the results of the synteny analysis with garlic. Not only the 3,123 onion EST markers showed a uniform distribution on the garlic and onion pseudomolecules (Fig 1), also the distribution of transposable elements is uniform over the garlic and onion chromosomes (Sun *et al.* 2020; Fig S7). Our hypothesis is in line with previous results in which the mapping of genes on physical chromosomes using molecular cytogenetic methods showed that the genes in Allium are localized in all three regions of the chromosome arms: proximal, interstitial and distal (Scholten *et al.* 2007; Masamura *et al.* 2012; Khrustaleva *et al.* 2016, 2019b, 2019a).

**Figure 1.**
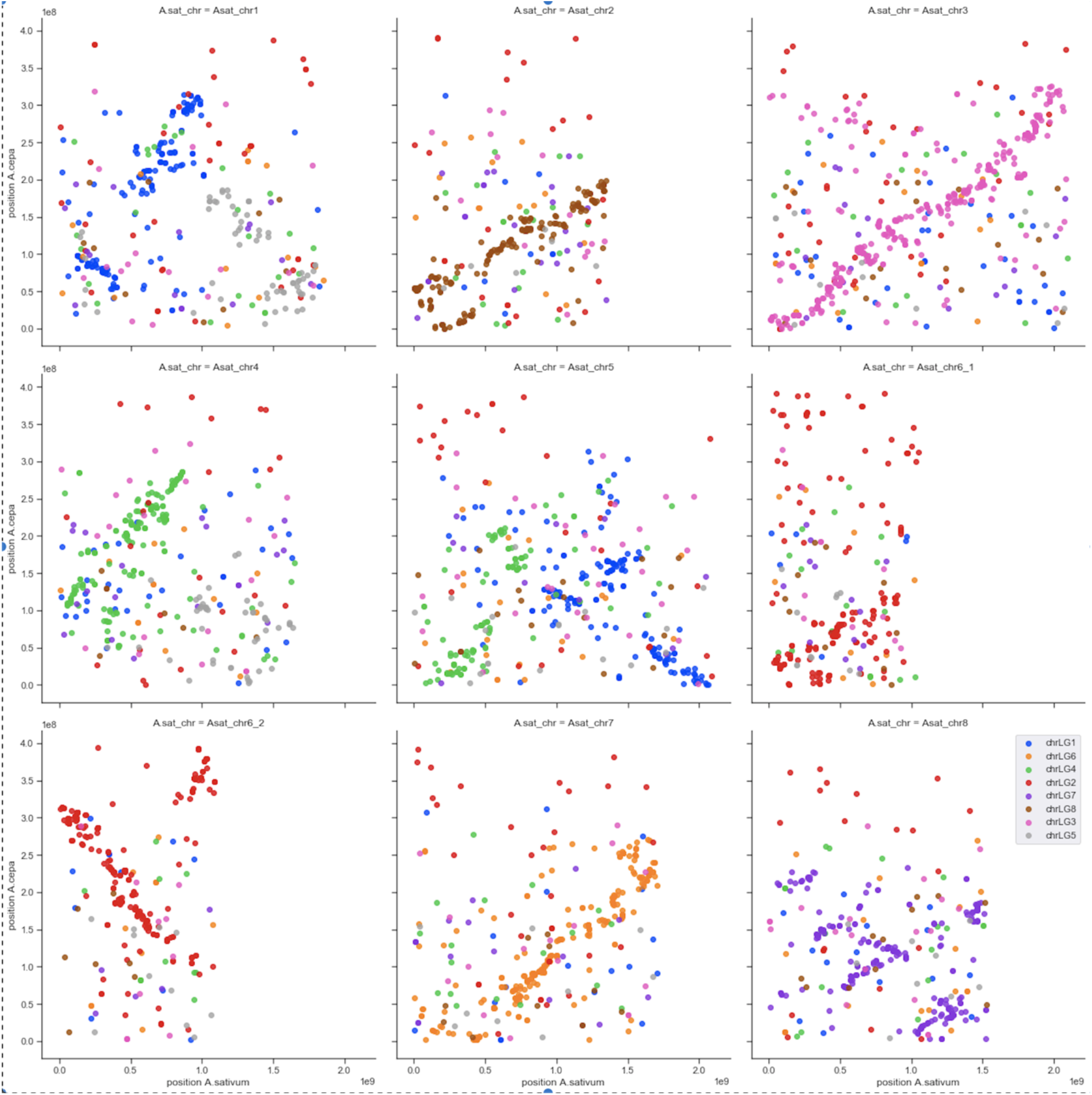
Physical positions of the marker on the garlic pseudomolecules (x-axis) plotted against the physical position of the marker on the onion pseudomolecules (y-axis). Color of the points represent the original linkage group assignment of the marker on the onion genetic map. Synteny between the garlic and onion genome can be observed in several chromosomes (e.g., chr3 and 7) though signals of translocation can also be observed (e.g., chr5). The data suggests inversions within a chromosome (e.g., chr8), but this cannot be estimated with certainty as not all our contigs could be oriented using Allmaps.

### Synteny between *A. cepa* (onion) and *A. sativum* (garlic)

Onion EST based markers (Shigyo 2020) were also used to study the synteny between onion and garlic (Sun *et al.* 2020). For 3,123 markers out of the 4,340, physical positions were determined on both the garlic and onion chromosomes. Overall synteny between some chromosomes is strong, such as for garlic chromosomes 3 and 7 and their onion counterparts chrLG3 and chrLG6 (Figure 1). Signals for translocations between chromosomes were also observed, as garlic chromosome 5 seems to be split over onion chrLG1 and chrLG4. In addition, signals for inversions were observed, for instance for chromosome 8. Where the signal for syntenic relationships between garlic and onion genomes are high, the garlic genome sequence may be used

### Improving usability of the onion genome assembly

The onion genome assembly in its current form is already a powerful tool for research and practical breeding. The annotation will be a good starting point for mining the genome for candidate genes while the set of pseudomolecules will facilitate the development of new markers for targeted regions. As syntenic relationships between garlic and onion genomes are high, insight in the overall synteny between garlic and onion can be used to develop hypothesis and assign unplaced scaffolds to approximate positions on the onion pseudomolecules, further facilitating discovery of novel insights. However, real improvements should come from additional lab work. Our current assembly is primarily based on Illumina short read sequencing and covers approx. 91% of the expected genome size and has a BUSCO completeness value 87.7% indicating, this assembly still needs improvement. Current third generation sequencing technologies, such as ONT long read (Jain *et al.* 2016) and PacBio HiFi (Wenger *et al.* 2019), have shown to deliver larger continues genome assemblies (Michael and VanBuren 2020). Data from one or both platforms, combined with, for example, Hi-C scaffolding data, would lead to a more continuous assembly, as shown for garlic (Sun *et al.* 2020), a genome with a size similar to onion.

### Conclusion

We have produced the first *de novo* genome sequence of onion. The sequence provides insights into the distribution of genes and repeats in this important cop species. This assembly is the first high-quality genome sequence and will be a valuable resource for both research and breeding.

## Acknowledgments

The authors would like to thank Dr Marta Mutschler for providing the DH line and Dr Alexey Zimin for his help in debugging the MaSurCa assembly pipeline. This work was, in part, carried out on the Dutch national e-infrastructure with the support of SURF Cooperative.

This research was supported by a grant from the Top Sector Horticulture & Propagation Materials (H279-SEQUON) and by the companies Bejo Zaden B.V., De Groot en Slot B.V., and GenomeScan. Within the Top Sector, the business community, knowledge institutions and the government work together on innovations in the field of sustainable production of healthy and safe food and the development of a healthy, green living environment.

## Supplementary information

**Table S1.** Summary Table of the generated sequencing data

| Lib | Tissue | DNA/<br>RNA | Nr of<br>Reads | Mean<br>Insert<br>Size<br>(bp) | Yield<br>(Gb) | Expected<br>Genome<br>Coverage |
| --- | --- | --- | --- | --- | --- | --- |
| **G1** | Leaf | DNA | 1,115M | 236 | 281 | 17.5 |
| **G2** | Leaf | DNA | 993M | 352 | 250 | 15.6 |
| **G3** | Leaf | DNA | 945M | 513 | 238 | 14.8 |
| **PB1\*** | Leaf | DNA | 11M | - | 116 | 7.3 |
| **R1** | Leaf | RNA | 24.6M | - | 4.9 | ND |
| **R2** | Flower | RNA | 19.5M | - | 3.9 | ND |
| **R3** | Root | RNA | 115.9M | - | 23.2 | ND |
| **R4** | Basal<br>plate | RNA | 14.2M | - | 2.8 | ND |

\*PacBio RSII, other data all with Illumina HiSeq

### File S2 DGB2OLC scaffolding

DBG2OLC (Ye *et al.* 2016b) was run with varying settings for kmerSize, KmerCov, MinOverlap and AdaptTH. Final DBG2OLC assembly was performed with KmerSize=21, KmerCov =2, MinOverlap=20 and AdaptThr=0.05 (run dg2olc_1000i). This setting was selected because of several criteria, including CEGMA (Parra *et al.* 2007) completeness, assembly size, number of Pacbio reads incorporated in the raw backbone (File S3).

**Figure S2.1.**
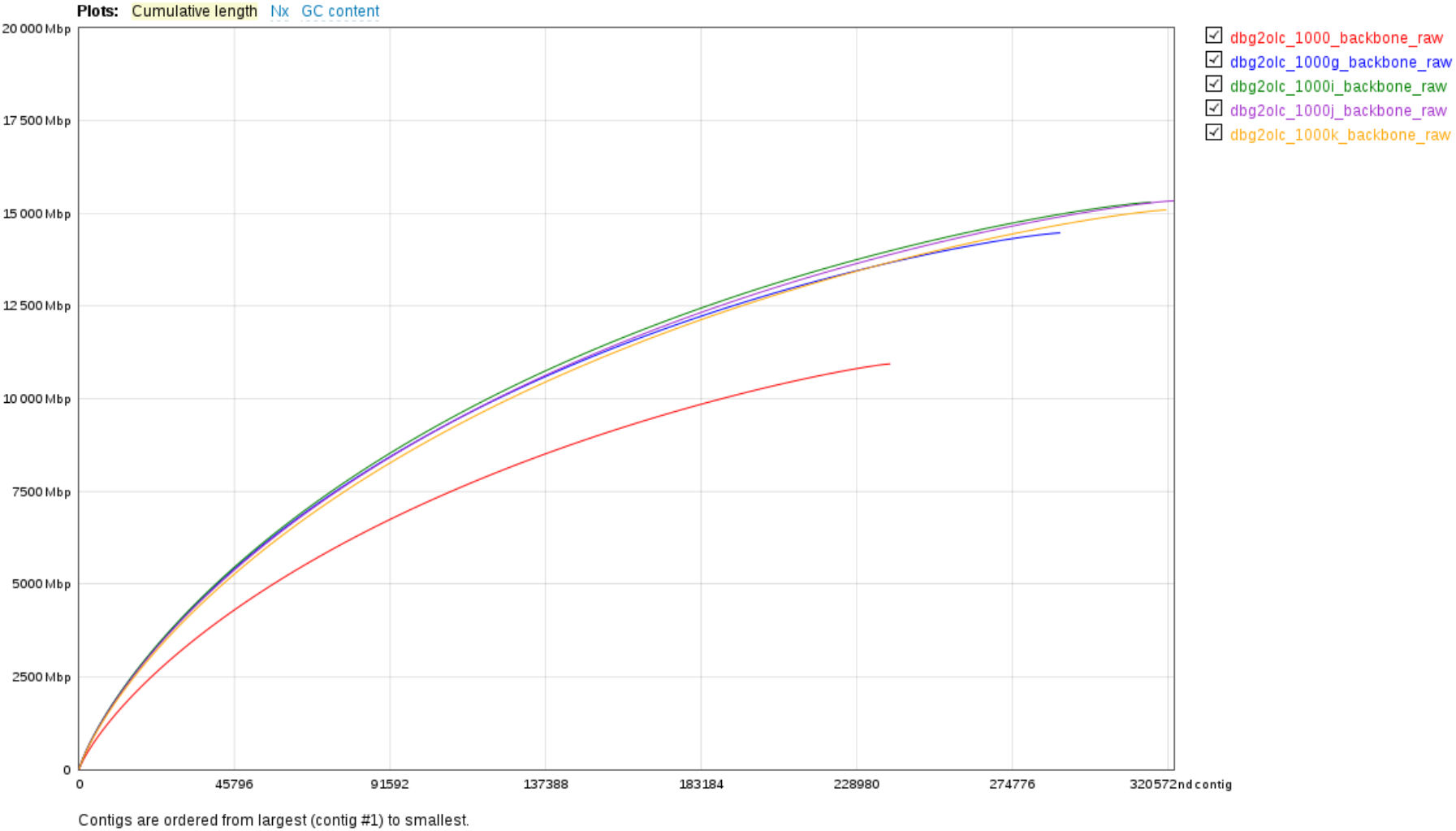
Quast quality control, showing total number of contigs and cumulative sequence length for four best performing dbg2olc runs and a baseline run with minor improvements (dbg2olc_1000_backbone_raw)

### File S4 – BUSCO Validation

\# BUSCO version is: 4.1.4

\# The lineage dataset is: embryophyta_odb10 (Creation date: 2020-09-10, number of species: 50, number of BUSCOs: 1614)

\# BUSCO was run in mode: genome

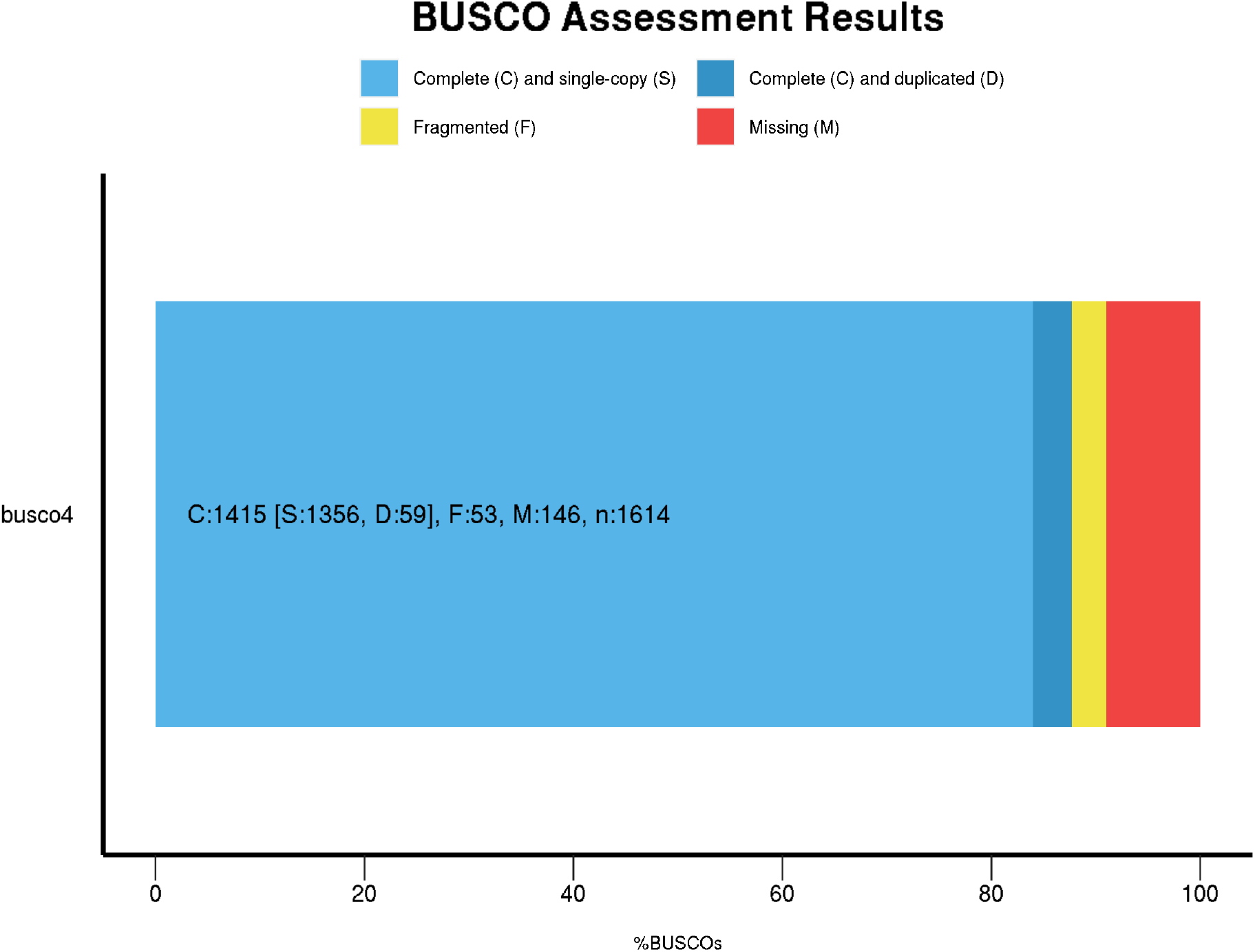

### Table S5 – Repeat Content and classification

Repeatmask

bases masked: 2258193827 bp or 2,25 Gb (15.12 %)

TEdenovo

bases masked: 10818118130 bp or 10.82 Gb (72.42 %)

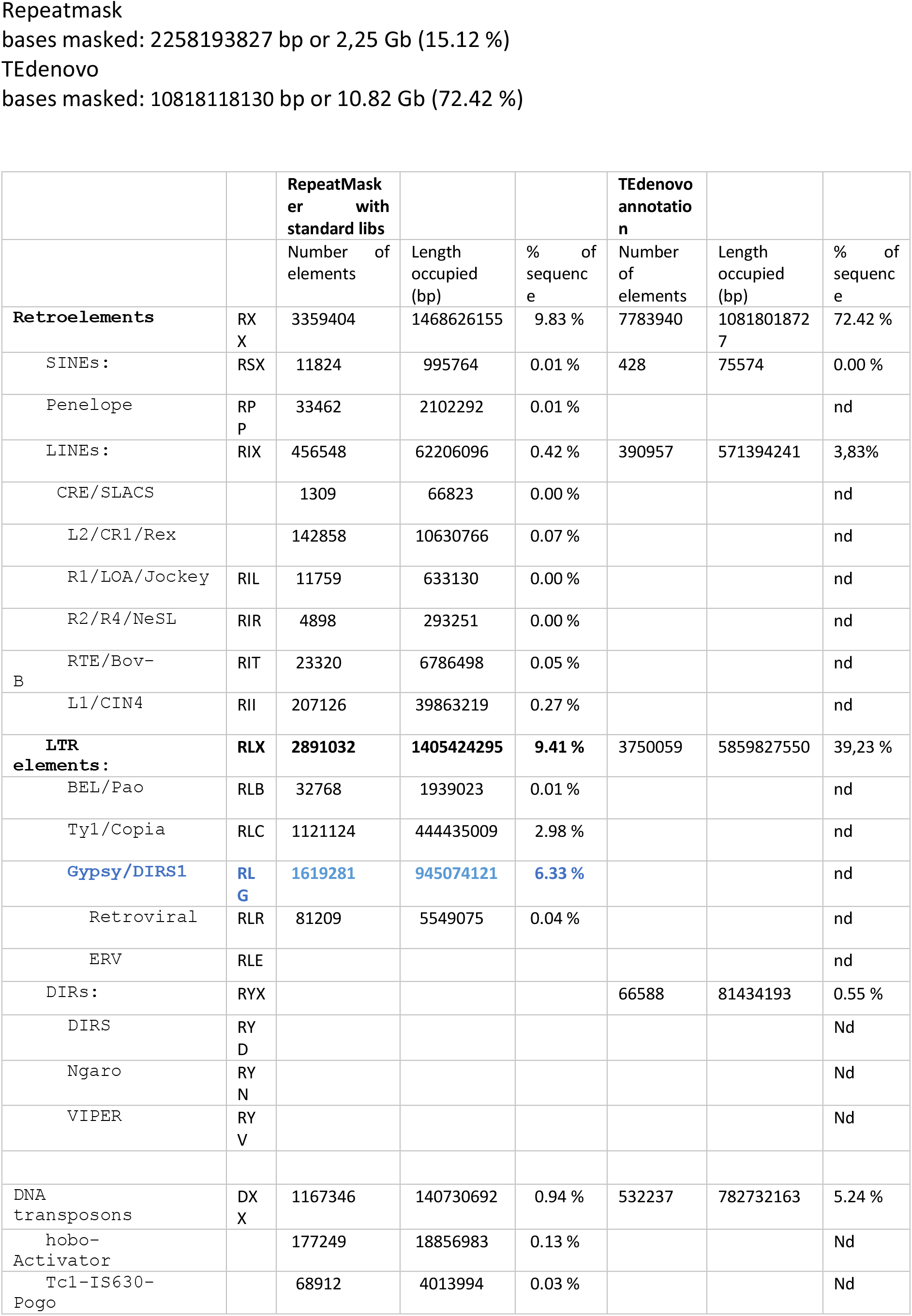
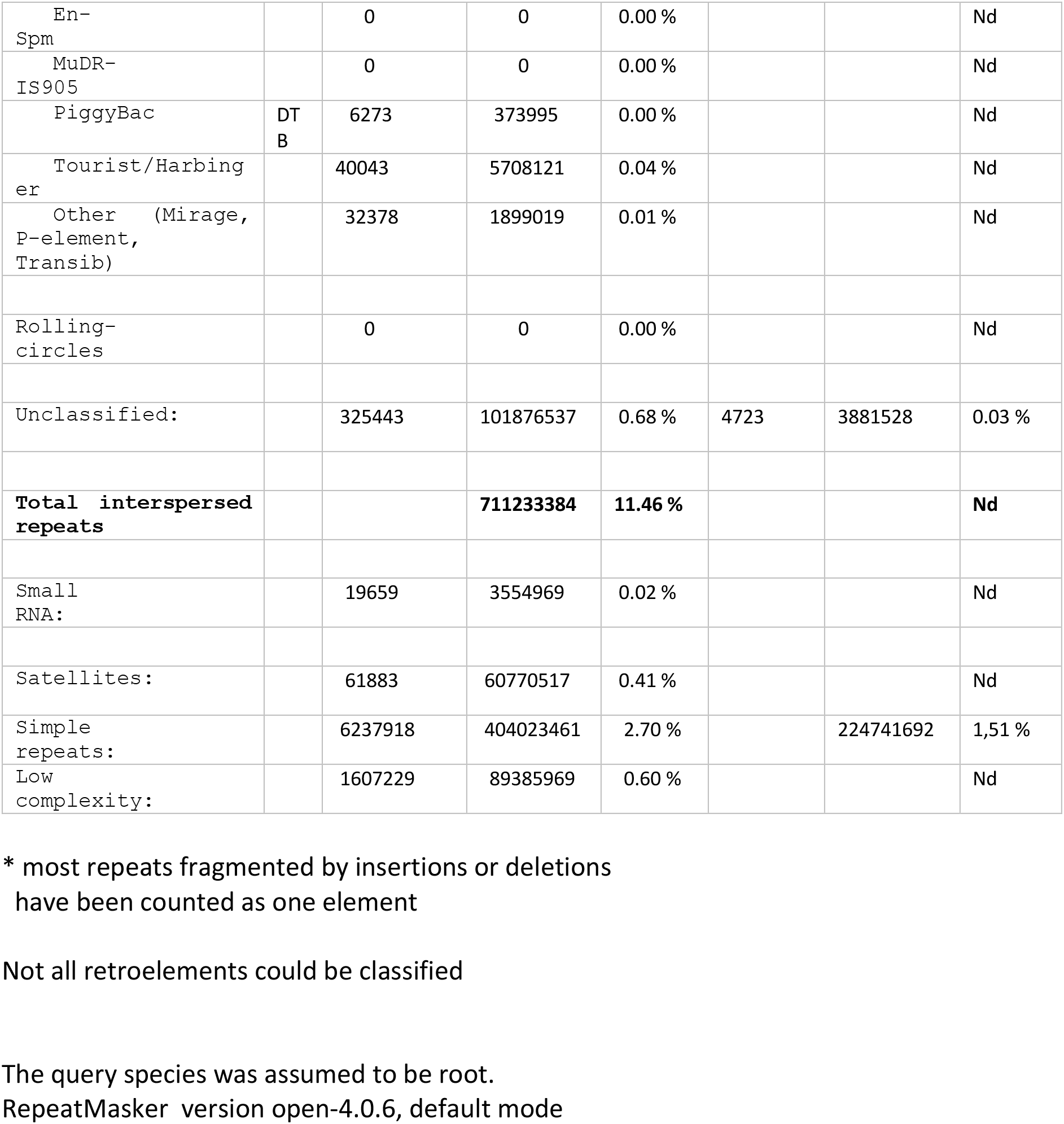

### File S6 – Genetic Anchoring Allmaps

Details of the Allmaps scaffolding (Tang *et al.* 2015) using five genetic linage maps. Scaffolding started with 5,815 markers, of which 4,926 were contributing to the final set of super scaffolds (Table S6.1). Furthermore, a subset of 1,747 markers was furthermore informative for orienting the scaffolds without conflicts given the five genetic maps. In total 2.2Gb (14.9%) of the overall assembly length could be scaffolded using this approach. This led to super scaffolds varying between 187Mb and 394 Mb in length (Table S6.2). Alignment of the individual genetic linkage maps to the final super scaffolds is shown in Figure S6.2. Overall correlation between a linkage group and its corresponding super scaffold is show in table S6.1. Correlation of specific maps is discussed in the main manuscript. The proposed set of super scaffolds is consolidated and made available as the first set of onion pseudomolecules.

**Figure S6.1.**
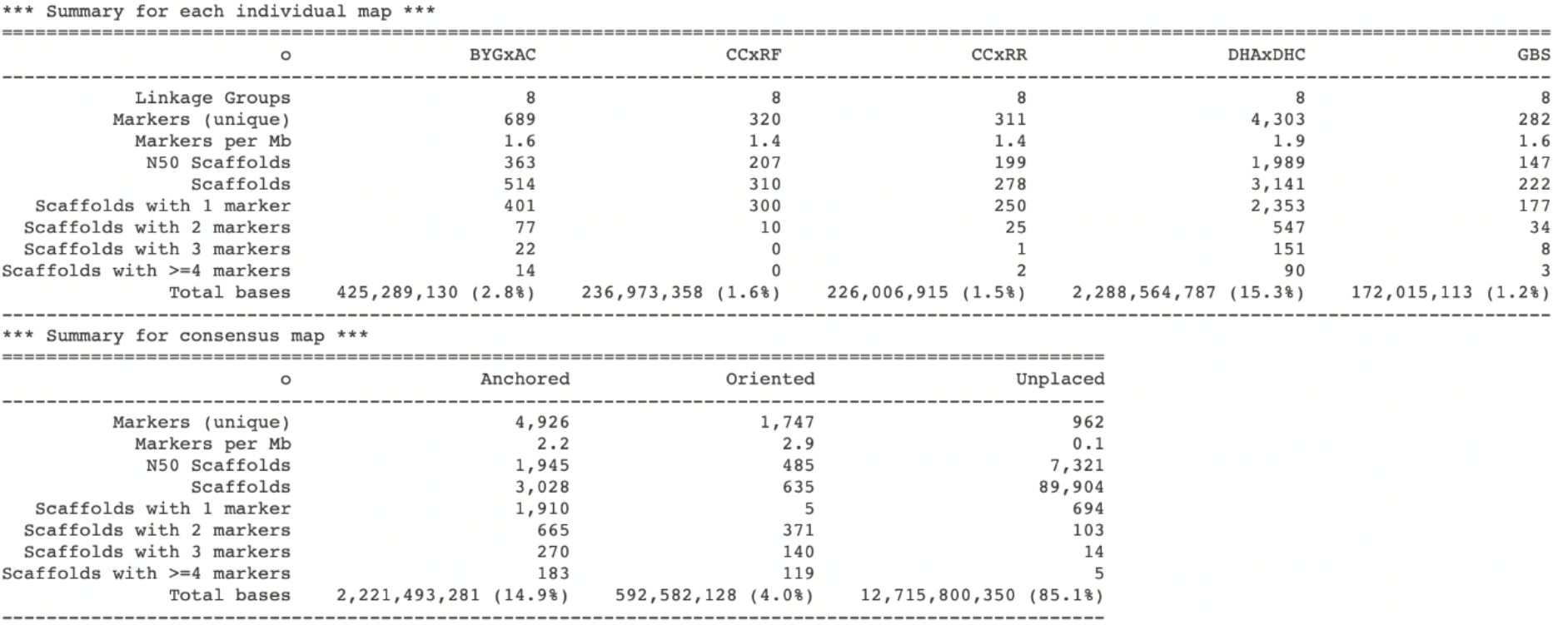
summary of the Allmaps scaffolding

**Table S6.1.** Spearman rho coefficients as a measure between the physical pseudomolecule representation and the individual marker order in the used genetic maps. The average spearman rho coefficient was also calculated as an absolute value over all linkage groups per map or per linkage group over all maps. Information from the interspecific maps is shown in italics. *No informative markers were identified by the Allmaps algorithm contributing to the overall scaffolding.

|  | BYGxAC | CCxRF | CCxRR | GBS | DHxHHC | Average |
| --- | --- | --- | --- | --- | --- | --- |
| LG1 | -0.941 | -0.945 | -0.909 | 1.000 | 0.960 | **0.951** |
| LG2 | 0.736 | -0.287 | 0.637 | 0.985 | -0.842 | **0.697** |
| LG3 | -0.840 | * | -0.501 | * | 0.871 | **0.737** |
| LG4 | 0.970 | 0.960 | 0.958 | -0.982 | -0.989 | **0.972** |
| LG5 | -0.856 | 0.873 | 0.836 | * | 0.917 | **0.871** |
| LG6 | 0.696 | 0.904 | -0.386 | 0.830 | 0.894 | **0.742** |
| LG7 | -0.733 | 0.541 | -0.316 | -0.943 | 0.915 | **0.690** |
| LG8 | 0.902 | 0.644 | 0.994 | * | -0.940 | **0.870** |
| **Average** | **0.834** | **0.736** | **0.692** | **0.948** | **0.916** |  |

**Table S6.2.**
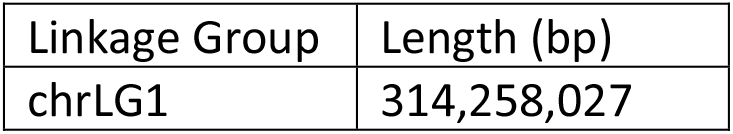
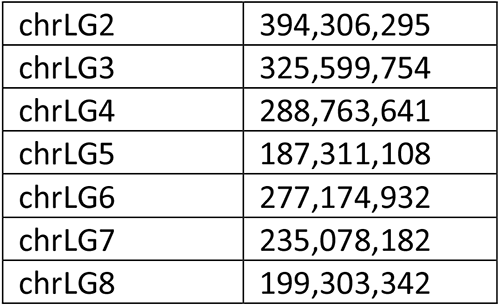
Size of the pseudomolecules after Allmaps scaffolding chrLG1

| Linkage Group | Length (bp) |
| --- | --- |
| chrLG1 | 314,258,027 |
| chrLG2 | 394,306,295 |
| chrLG3 | 325,599,754 |
| chrLG4 | 288,763,641 |
| chrLG5 | 187,311,108 |
| chrLG6 | 277,174,932 |
| chrLG7 | 235,078,182 |
| chrLG8 | 199,303,342 |

**Figure S6.2.**
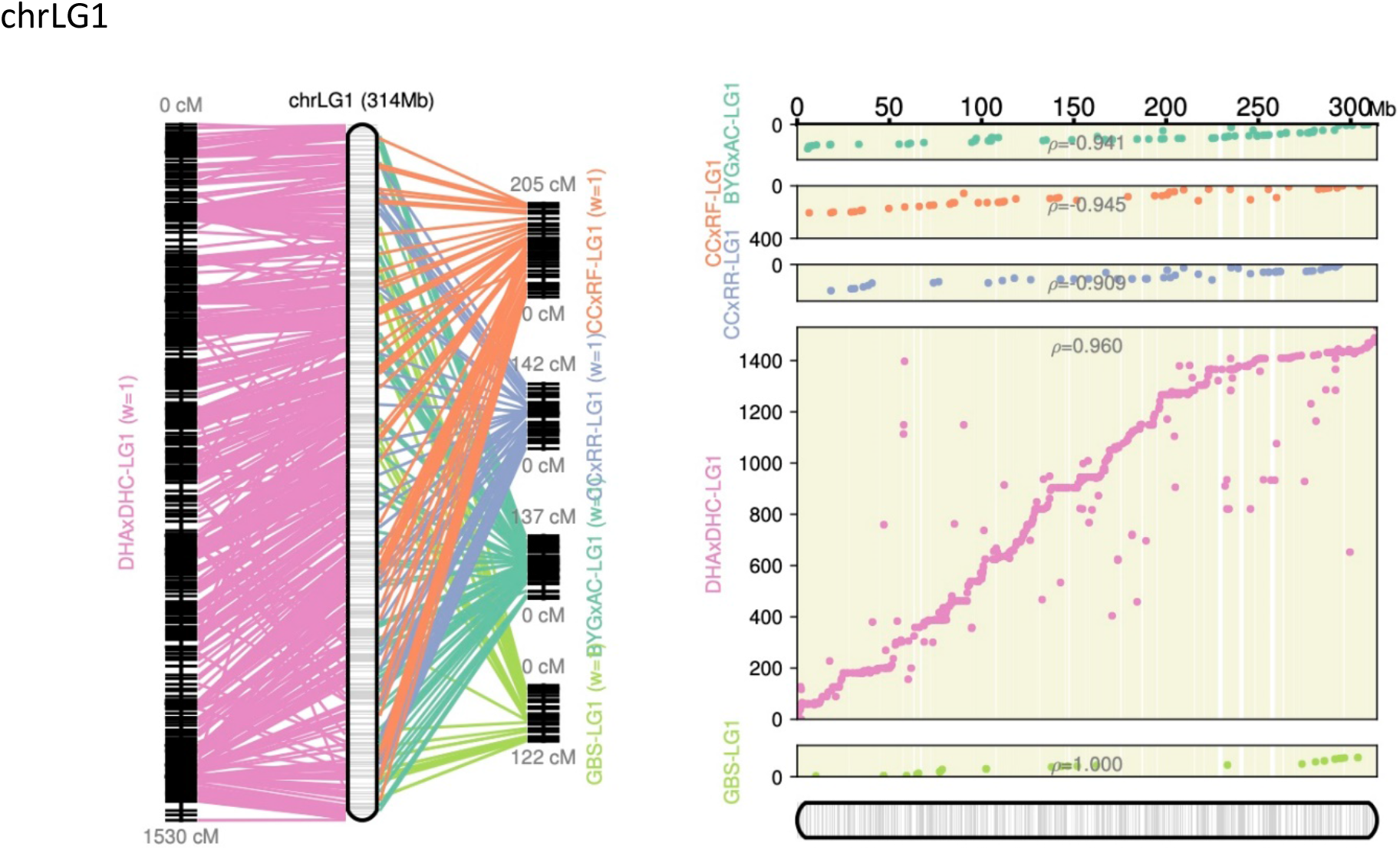
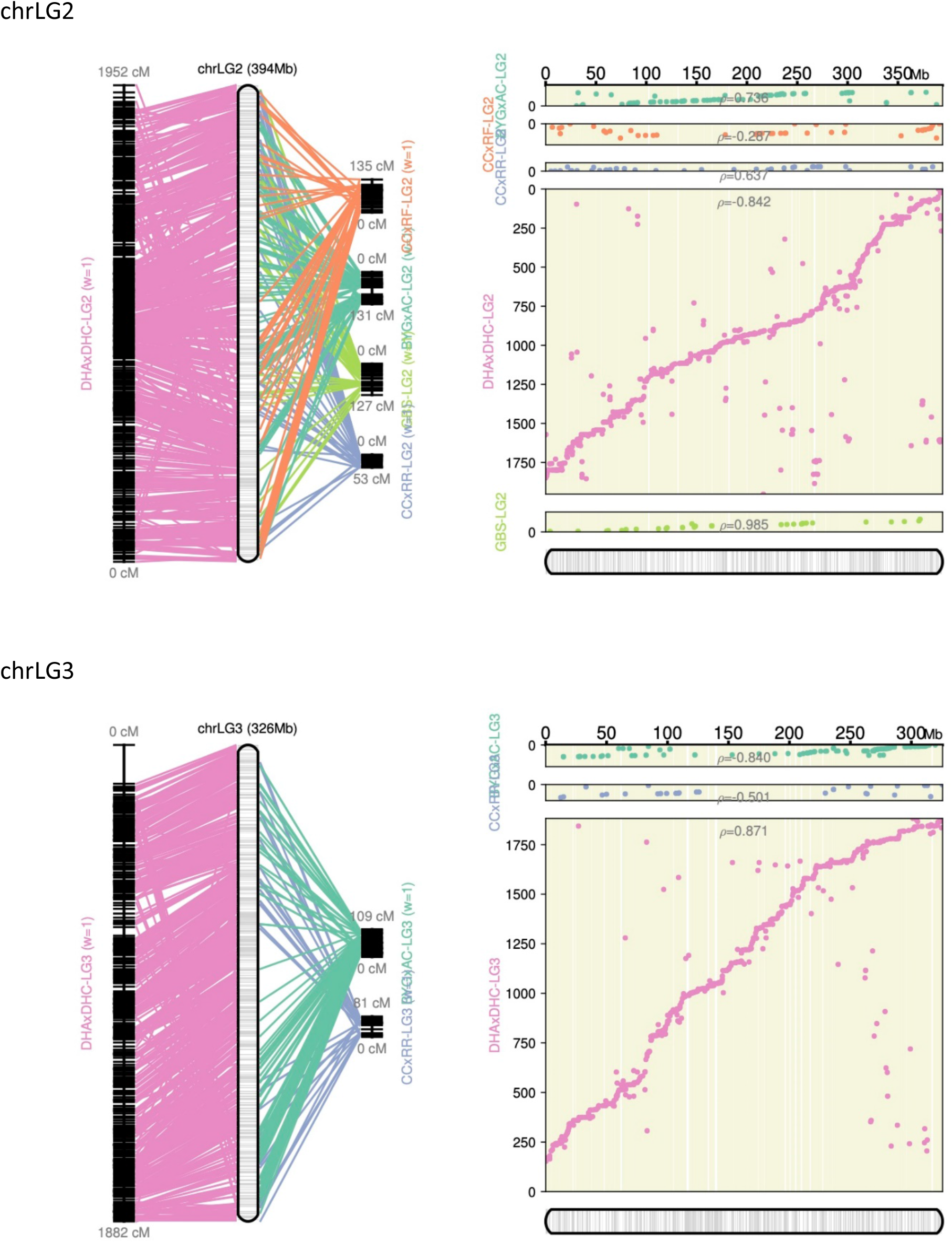
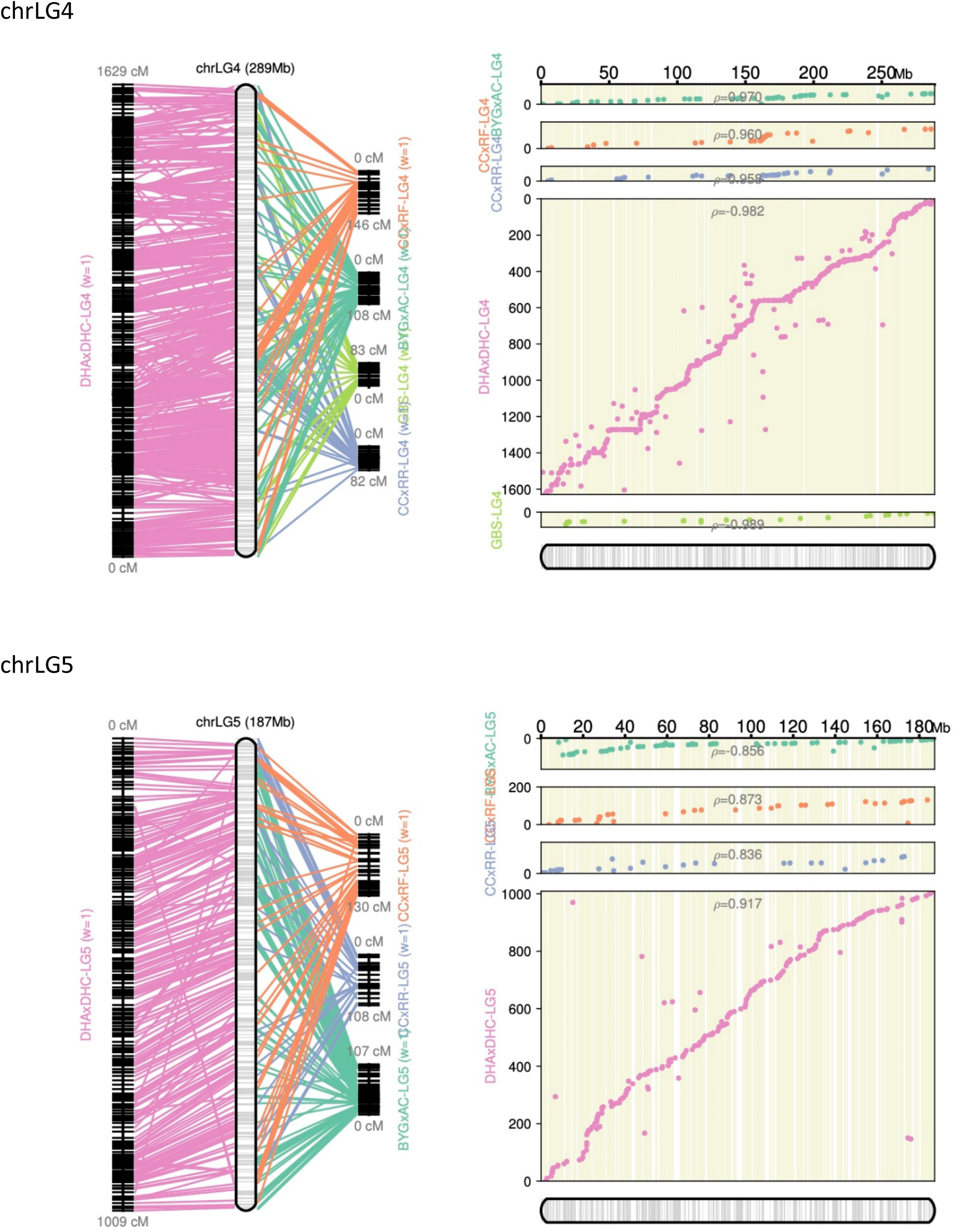
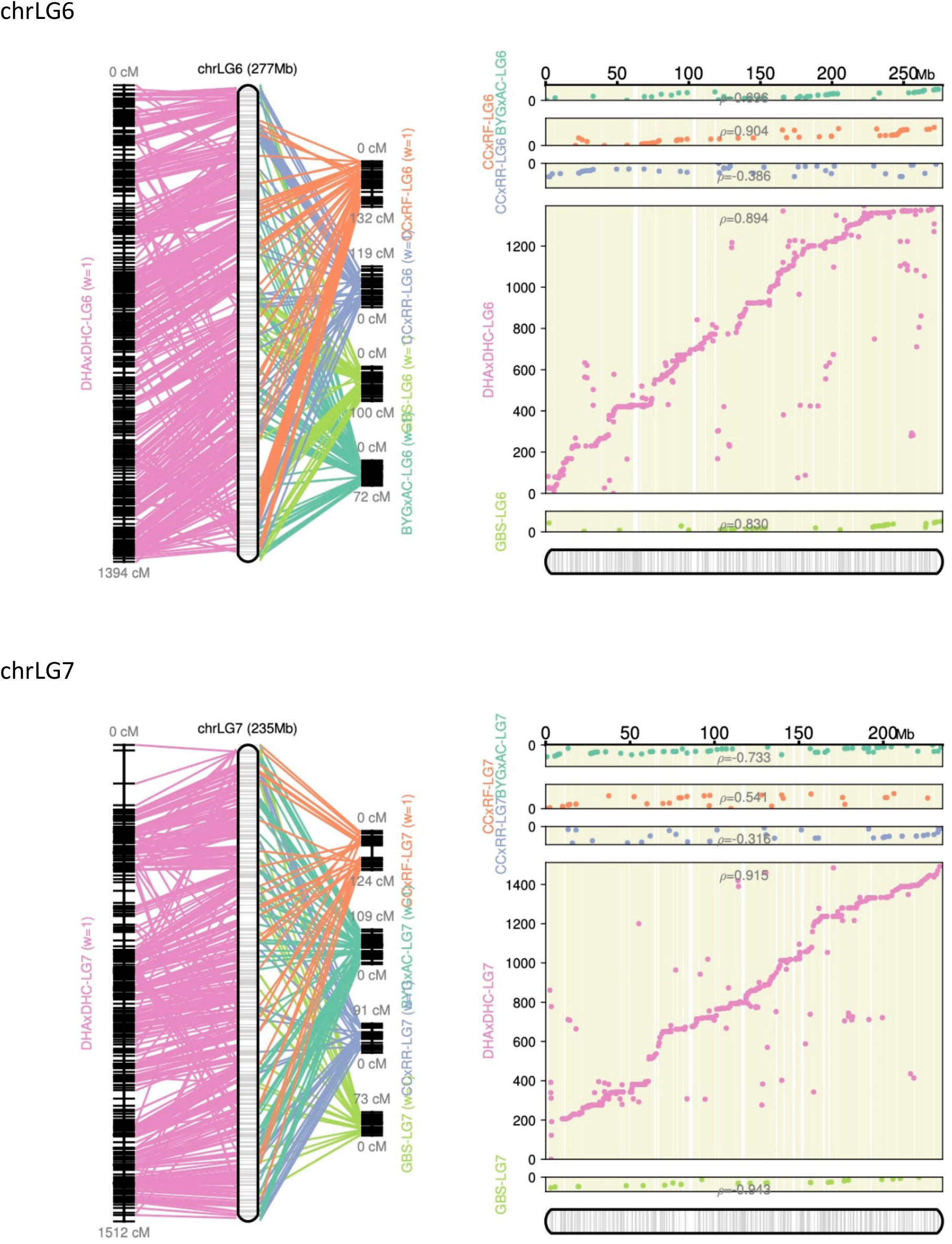
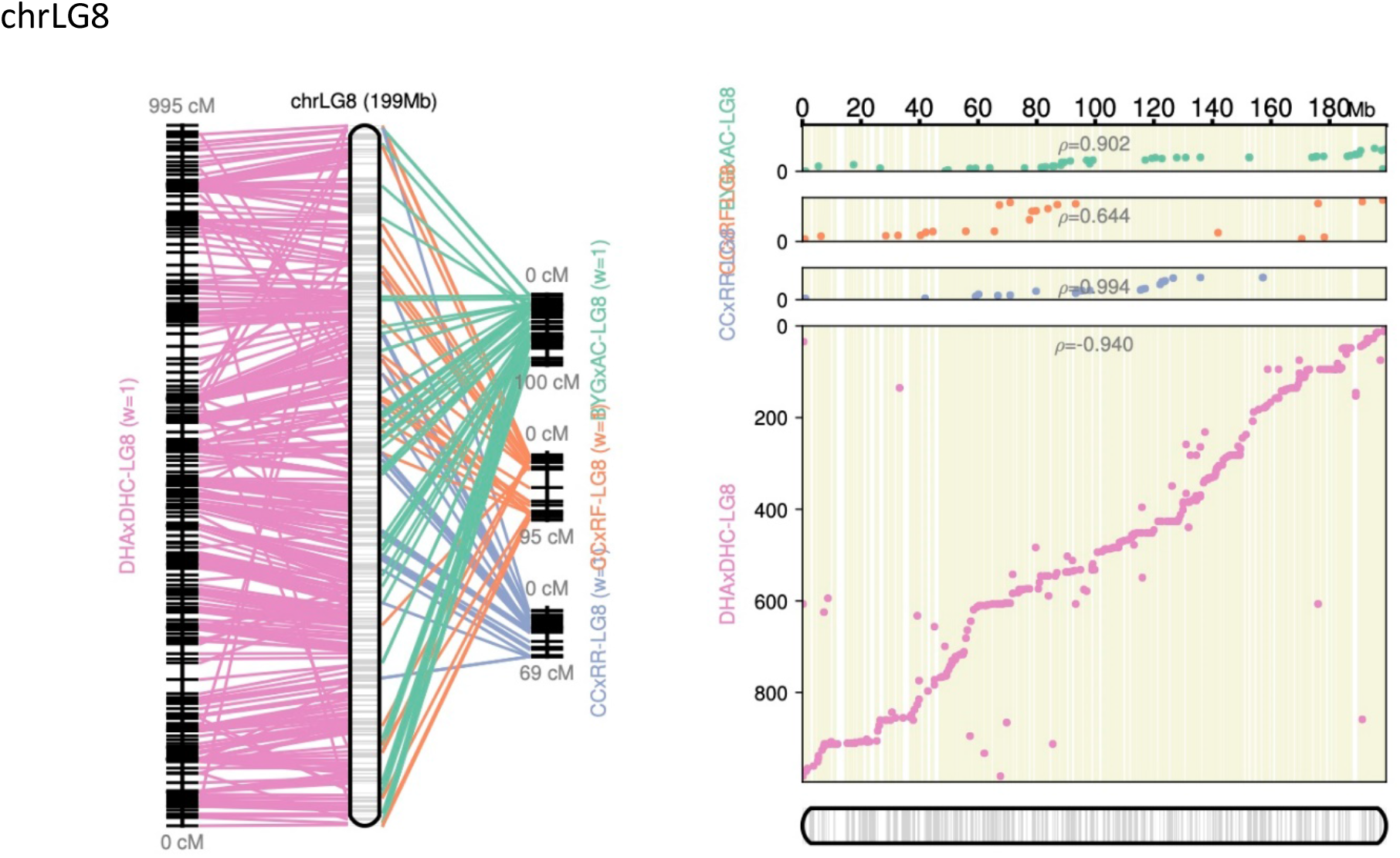
Alignment of the genetic linkage maps to the final ordering of the contigs into pseudomolecules.

### Figure S7 – Gene and repeat landscape of the pseudomolecules

**Figure S7.1.**
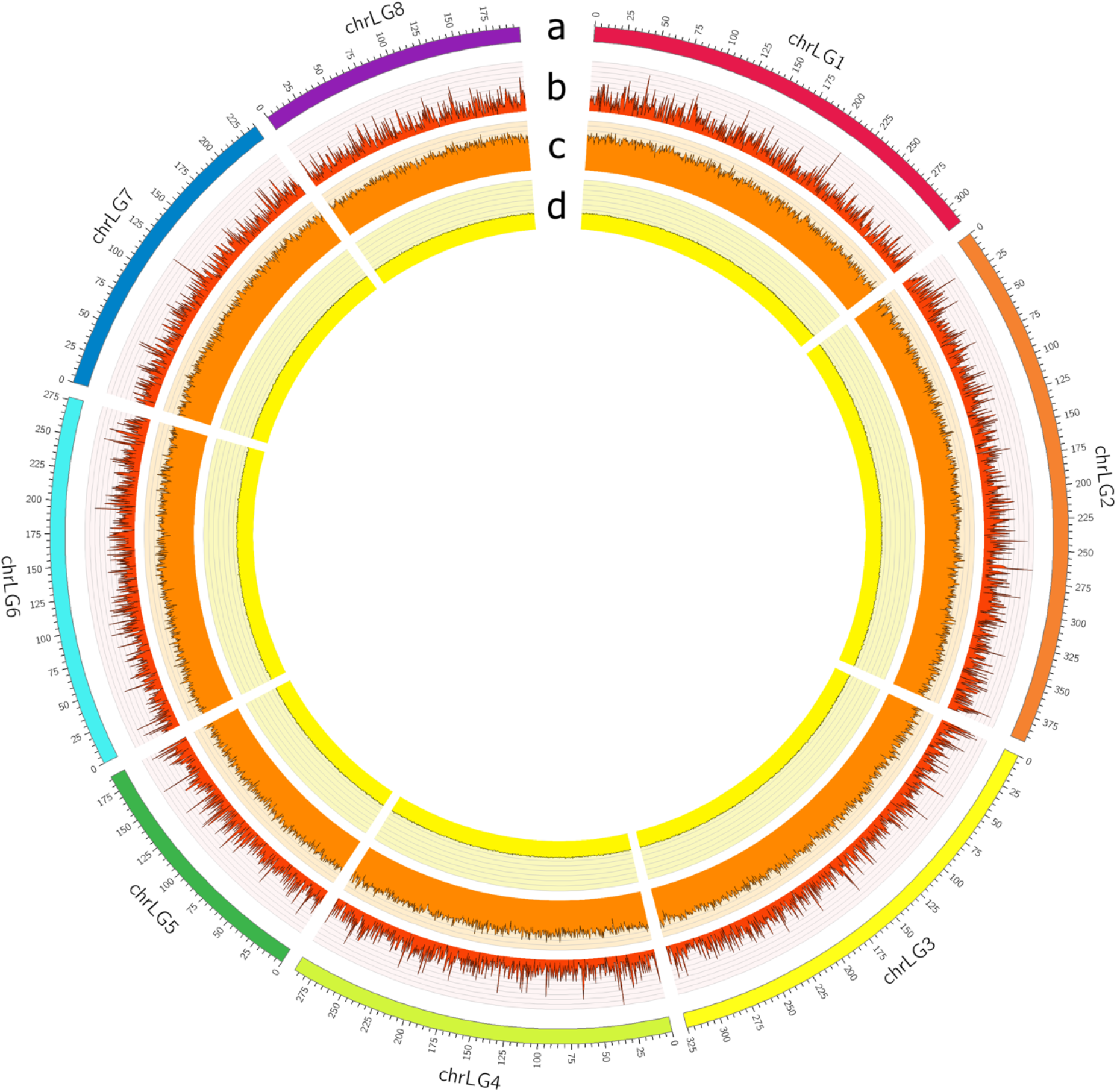
Circos plot showing a) pseudomolecule length, b) gene density per pseudomolecule (binsize 500kb), c) transposable element (TE) content per pseudomolecule (binsize 500kb), and d) distribution of GC content per pseudomolecule (binsize 500kb)

